# Synthetic glycans that control gut microbiome structure mitigate colitis in mice

**DOI:** 10.1101/2022.01.28.478269

**Authors:** Andrew C Tolonen, Nicholas Beauchemin, Charlie Bayne, Lingyao Li, Jie Tan, Jackson Lee, Brian Meehan, Jeffrey Meisner, Yves Millet, Gabrielle LeBlanc, Robert Kottler, Erdmann Rapp, Chris Murphy, Peter J Turnbaugh, Geoffrey von Maltzahn, Christopher M Liu, Johan ET van Hylckama Vlieg

## Abstract

Relative abundances of bacterial species in the gut microbiome have been linked to many diseases. Species of gut bacteria are ecologically differentiated by their abilities to metabolize different glycans, making glycan delivery a powerful way to alter the microbiome to promote health. We describe the properties and therapeutic potential of chemically diverse synthetic glycans (SGs). Fermentation of SGs by gut microbiome cultures resulted in compound-specific shifts in taxonomic and metabolite profiles not observed with reference glycans, including prebiotics. Model enteric pathogens grow poorly on most SGs, potentially increasing their safety for at-risk populations. SGs increased survival, reduced weight loss, and improved clinical scores in mouse models of colitis. Synthetic glycans are thus a promising modality to improve health through selective changes to the gut microbiome.

## Introduction

Gut microbiome composition and metabolic output have been associated with initiation and progression of diseases including auto-immune diseases, cancer, metabolic syndrome, and liver disease^1-5^. Targeted manipulation of the microbiome is thus a promising therapeutic strategy being pursued by numerous public and private initiatives to treat disease^6^. In particular, glycan intervention is a chemical-based approach to alter the microbiome that leverages how ecological niches in the gut are occupied by species that are specialized to metabolize different carbon sources^7,8^. Alimentary interventions with complex glycans comprising dietary fiber significantly shift the composition and output of the gut microbiome within a few days and have been linked to prevention of type 2 diabetes, obesity, and cancer^9-12^.

Glycans with documented health benefits currently marketed as prebiotics include fructo-oligosaccharides (FOS), galacto-oligosaccharides (GOS), xylo-oligosaccharides (XOS), pullulan, and lactulose^13-17^. These glycans are extracted from agricultural materials or enzymatically synthesized and lack the structural complexity of dietary fiber, which is composed of a matrix of β-glucans, starches, hemicelluloses, and pectins with diverse monosaccharides and glycosidic bonds. To exploit the potential of glycans to modulate the microbiome, there is a need for novel molecules spanning the chemical and structural diversity of dietary glycans that can be efficiently and consistently produced. By controlling the starting materials and reaction conditions, we synthesized a library of hundreds of synthetic glycans (SGs) with different monosaccharides, bond types, and degrees of polymerization (DPs)^18, 19^. These SGs enable a wide range of targeted changes to the microbiome and potentially open new avenues for the prevention and treatment of disease.

In this study, the fermentations of hundreds of SGs polymerized from diverse, naturally occurring monosaccharides were compared to reference glycans using an *ex vivo* platform for highly multiplexed measurements of growth, metabolic output, and taxonomy. Based on the *ex vivo* assays, we selected SGs for chemical structural analyses of their polymerization profiles and glycosidic linkages. As some reference glycans have been shown to alleviate intestinal inflammation and improve barrier function, as well as reduce colitis from infection, we compared the effects of SG and reference glycan treatment in mouse models of these intestinal pathologies^20, 21^. Together, this experimental pipeline (Fig 1A) evaluated the function and therapeutic potential of SGs as microbiome modulators and demonstrated additional benefits of SGs compared to reference glycans, including compounds currently marketed as prebiotics.

**Fig 1.**
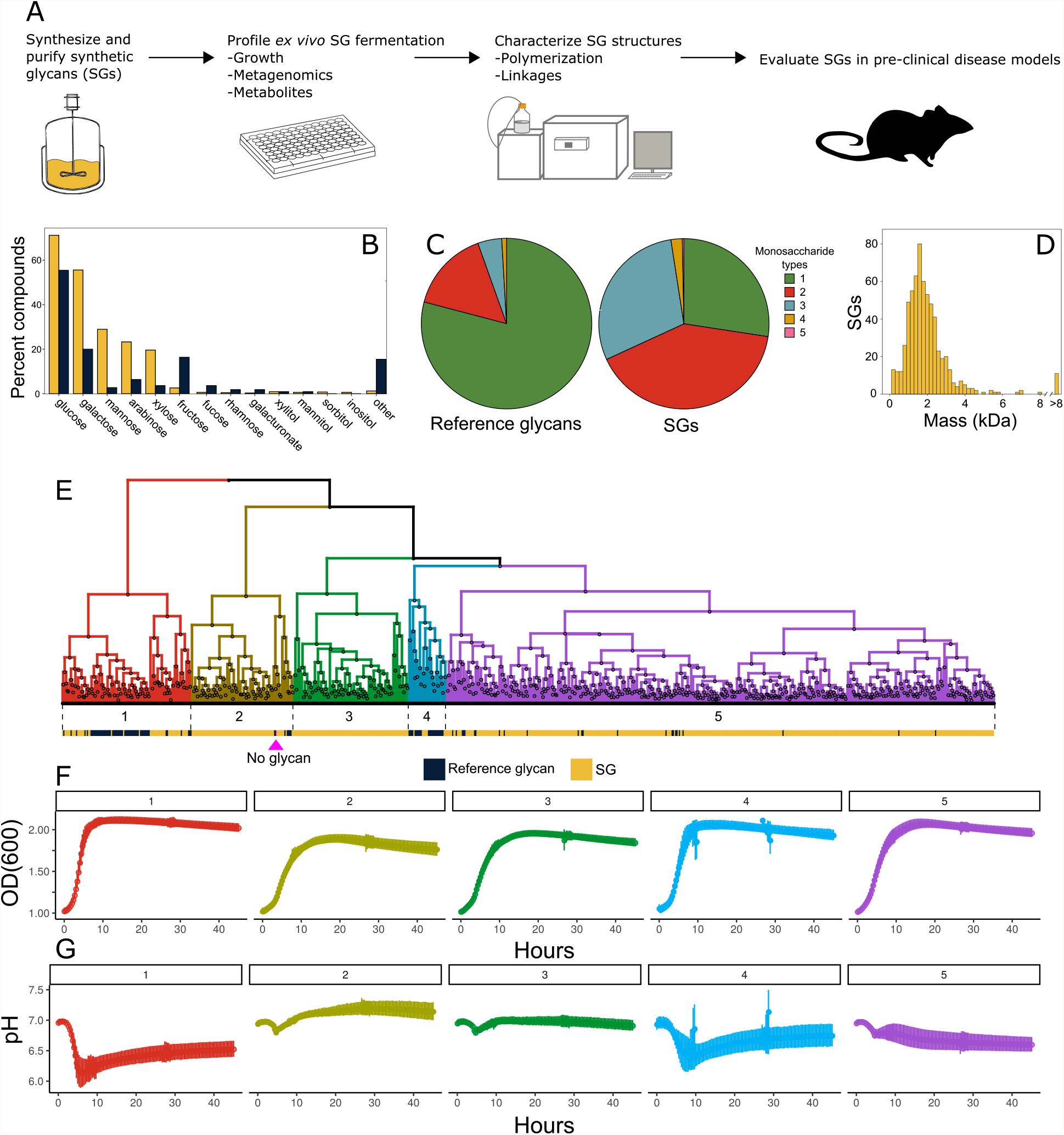
(A) Schematic representation of the analytical pipeline. (B-F) Compositions and fermentation dynamics of a set of 653 SGs and 110 reference glycans. (B) Percentages of SGs (yellow) and reference glycans (indigo) containing various monosaccharide types. (C) Number of monosaccharide types composing each SG or reference glycan. (D) Distribution of weight-average molecular weights of SGs measured by SEC. (E-G) Growth (OD_600_) and pH dynamics of triplicate fecal cultures fermenting 5 g l^-1^ of a single SG or reference glycan in MM29 medium. (E) Hierarchical clustering of glycans into five fermentation groups based on twelve growth and pH parameters. Bars below the dendrogram show compound class: SG (yellow), reference glycan (indigo) or no glycan (magenta). Mean (F) growth and (G) pH curves (±SD) for each glycan fermentation group shown in (E). SGs, Synthetic Glycans; SEC, size exclusion chromatography; kDa, kilodalton; OD_600_, optical density at 600 nm; SD, standard deviation.

## Results

### SGs vary in composition and fermentation dynamics

We selected a set of 653 SGs and 110 commercially available, reference glycans that are compositionally diverse and sufficiently soluble for culture-based growth and metabolite assays (Supplementary Table 1A). Both SGs and reference glycans are composed of a similar set of monosaccharides (Fig 1B), but unlike the reference glycans, most SGs (73%) contain multiple, different monosaccharides (Fig 1C, Supplementary Table 1B), demonstrating how SGs can be built to include dietary sugars in novel and complex combinations. SGs span a wide range of average molecular masses (Fig 1D) with a median of 1.7 kDa (range 0.3 kDa-77.5 kDa), corresponding to a polymerization of approximately 10 monosaccharides. We profiled these SGs and reference glycans in a panel of *ex vivo* assays and highlighted the performance of two SGs with different compositions, BRF (glucose) and BQM (galactose and glucose), which were subsequently selected for structural analysis and mouse models.

We investigated variation in fermentation dynamics across the 763 glycans by measuring growth and pH kinetics of anaerobic fecal cultures. Hierarchical clustering of growth and pH parameters (Fig 1E, Supplementary Fig 1, Supplementary Table 1A) identified groups of glycans with distinct growth (Fig 1F) and pH (Fig 1G) profiles. Reference glycans were enriched in group 1 (Fisher exact test p=7.5×10^−26^) and group 4 (Fisher exact test p=4.8×10^−14^), which both supported rapid growth resulting in a precipitous pH drop. Group 1 included pullulan, lactulose, GOS, and FOS; group 4 included XOS. Group 5, which included BQM and BRF, was enriched in SGs (p= 6.0×10^−17^) that were well-fermented at controlled rates with gradual reductions in pH. Thus, SG composition affects fermentability and SGs are generally fermented more slowly than reference glycans, likely due to their compositional complexity (Fig 1C). In addition, these fermentation parameters were well correlated between fecal samples from two different donors (Supplementary Fig 1C-N). While more data is needed to establish that SG effects are conserved across populations, these data suggest SGs have similar fermentation dynamics between individuals.

### SGs change microbiome metabolic output

Fermentation of glycans by the gut microbiome produces metabolites with important physiological and immunological benefits including short chain fatty acids (SCFAs)^15, 22^. We thus quantified butyrate and propionate yields in fecal cultures fermenting the 763 glycans. Butyrate is the preferred energy source of colonocytes, leading to healthy colonocyte function and maintenance of an anaerobic gut environment^23^. Propionate has immunological and metabolic effects locally in the gut and is absorbed into the bloodstream to regulate cholesterol^24, 25^. In addition, butyrate and propionate promote differentiation of naive T cells into anti-inflammatory regulatory T cells^26, 27^.

Butyrate and propionate production by some SGs were similar to those of reference glycans, but many SGs were distinct in producing high propionate levels (Fig 2A), supporting these SGs favor growth of propionate-producing taxa such as *Bacteroidaceae* and *Ruminococcaceae*^25^. Additionally, we investigated the extent to which SCFA production underlies pH dynamics observed during glycan fermentation by measuring a time series of total SCFA production for a subset of glycans. While the correlation between pH and SCFA production is weak, the relative levels of SCFA and ammonia, an abundant microbial product, were strongly correlated with PH (Supplementary Fig 2). Thus, changes in pH observed during glycan fermentation are associated with the balance between SCFA and ammonia production.

**Fig 2.**
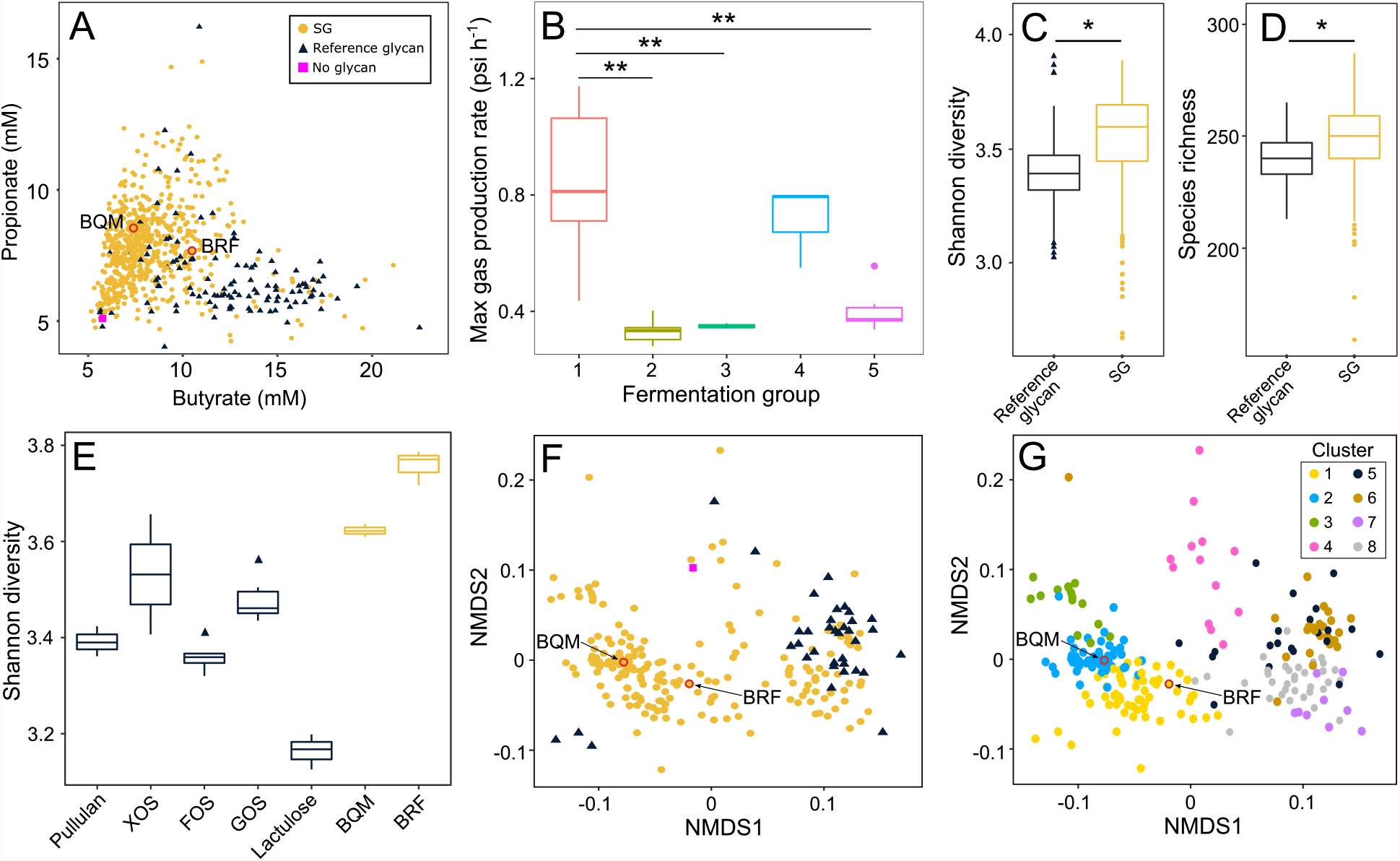
Effects of glycans on fecal community (A-B) metabolic output and (C-G) taxonomic composition based on metagenomics. (A) Yields of two SCFAs, butyrate and propionate, from fecal cultures fermenting either an SG (yellow circles, n=653), reference glycan (indigo triangles, n=110), or no glycan (magenta square). (B) Maximum gas production rate (psi h^-1^) during fecal culture fermentation of glycans from each of the five fermentation groups in Fig 1E-G. (C) Shannon diversity and (D) species richness in fecal cultures fermenting SGs (yellow, n=190) versus reference glycans (indigo, n=40). (E) Shannon diversity of fecal cultures fermenting BRF or BQM (yellow) is higher than reference glycans (indigo) for all pairwise comparisons except BQM versus XOS (Tukey’s test: p<0.05). (F) NMDS of metagenomic data calculated based on a matrix of Bray-Curtis dissimilarities using species-level mapping of sequencing reads from fecal cultures grown on either an SG (yellow circles, n=190), reference glycan (indigo triangles, n=40), or no glycan (magenta square). (G) NMDS as in (F) colored by differences in taxonomic composition defined by eight K-means clusters based on species-level mapping of sequencing reads. Data shows means of (A-D, F, G) three or (E) six replicate fecal cultures grown on 5 g l^-1^ of each SG or reference glycan for 45 hours in MM29 medium. (A, F, G) BRF and BQM highlighted in red. (B-E) Box plots show median and interquartile range. Asterisks show significance (*p<0.05, **p<0.01) by Tukey’s test. SG, Synthetic Glycan; SCFA, short-chain fatty acid; FOS, fructo-oligosaccharides; GOS, galacto-oligosaccharides; XOS, xylo-oligosaccharides, NMDS, non-metric multidimensional scaling.

In addition to SCFAs, glycan fermentation produces gaseous compounds that can lead to abdominal bloating, a major symptom limiting tolerability of reference glycans in patients with gastrointestinal disorders such as irritable bowel syndrome^28^. As the volume and rate of gas produced depends on diet and microbiome composition^29^, we measured gas production rates by fecal communities growing on BRF, BQM, and randomly selected glycans from each of the five fermentation groups in Fig 1E-G (Supplementary Table 2). The rate of gas production varied widely among groups (Fig 2B) and reflected fermentation dynamics. Glycans in fermentation groups 1 and 4, mostly reference glycans that are rapidly fermented, had the most rapid gas production. SGs in group 5, which included BRF and BQM, produced gas more moderately, potentially improving tolerability in humans.

### SGs shift microbiome taxonomic composition

Glycan fermentation can result in divergent changes to microbiome composition, even differentially promoting closely-related species^8^. Therefore, we applied shotgun metagenomic sequencing to examine fecal cultures fermenting a compositionally diverse set of 190 SGs and 40 reference glycans (Supplementary Table 3). While there is heterogeneity in the level of microbiome diversity resulting from fermentation of different SGs, SG fermentation generally resulted in higher taxonomic diversity (Fig 2C) and species richness (Fig 2D) relative to reference glycans, as shown by BQM and BRF promoting higher diversity relative to reference glycans (Fig 2E).

Fecal community compositions following SG fermentation spanned the taxonomic space covered by reference glycan fermentation, but also reached novel compositions (Fig 2F) that can be explored by clustering based on taxa abundances. Clusters enriched in reference glycan fermentations contained elevated levels of *Lactobacillaceae* (cluster 5) and *Bifidobacterium* (clusters 6-8) (Fig 2G, Supplementary Fig 3E-H), which are adapted to metabolize simple oligosaccharides such as FOS^30^. SG-enriched taxa clusters had elevated abundances of *Lachnospiraceae* (clusters 1-3) and *Parabacteroides* (clusters 2-3) (Fig 2G, Supplementary Fig 3A-C). BRF is in glycan cluster 1 (Fig 2G, Supplementary Fig 3A) with increased abundances of members of the family *Lachnospiraceae* including *Roseburia*, a key butyrate producer that is reduced in the microbiomes of patients with Crohn’s disease^31^, and *Fusicatenibacter*, which is decreased in patients with active ulcerative colitis^32^. BQM is in glycan cluster 2 (Fig 2G, Supplementary Fig 3B) with elevated levels of other *Clostridiaceae* and *Parabacteroides*, a taxon associated with remission in Crohn’s disease^33^.

### SGs are poor growth substrates for model pathogens

It is important to establish that microbiome-modulating glycans do not promote proliferation of pathogens. In particular, intestinal colonization by a multidrug resistant (MDR) pathogen such as carbapenem-resistant *Enterobacteriaceae* (CRE) expressing extended-spectrum carbapenemases or vancomycin-resistant *Enterococcus* (VRE) is a major risk factor for bloodstream infection^34, 35^. Currently, no approved therapies exist for de-colonization of the gut from these pathogens. We hypothesized that SGs will confer a competitive growth advantage to polysaccharide-utilizing commensals relative to pathogens in gut microbiomes^36^. Initially, we examined the ability of a panel of laboratory and MDR clinical *Enterobacteriaceae* and *Enterococcus* strains (Supplementary Table 4A) to grow on each of a compositionally diverse set of 148 SGs including BRF and BQM and 32 reference glycans including FOS (Supplementary Table 4B) as a sole carbohydrate source in defined medium. Most strains of the model enteric pathogens *Klebsiella pneumoniae* (Fig 3A), *Escherichia coli* (Fig 3B), and *Enterococcus faecium* (Fig 3C) exhibited limited growth on SGs with final cell densities significantly lower than when grown on reference glycans including FOS.

**Fig 3.**
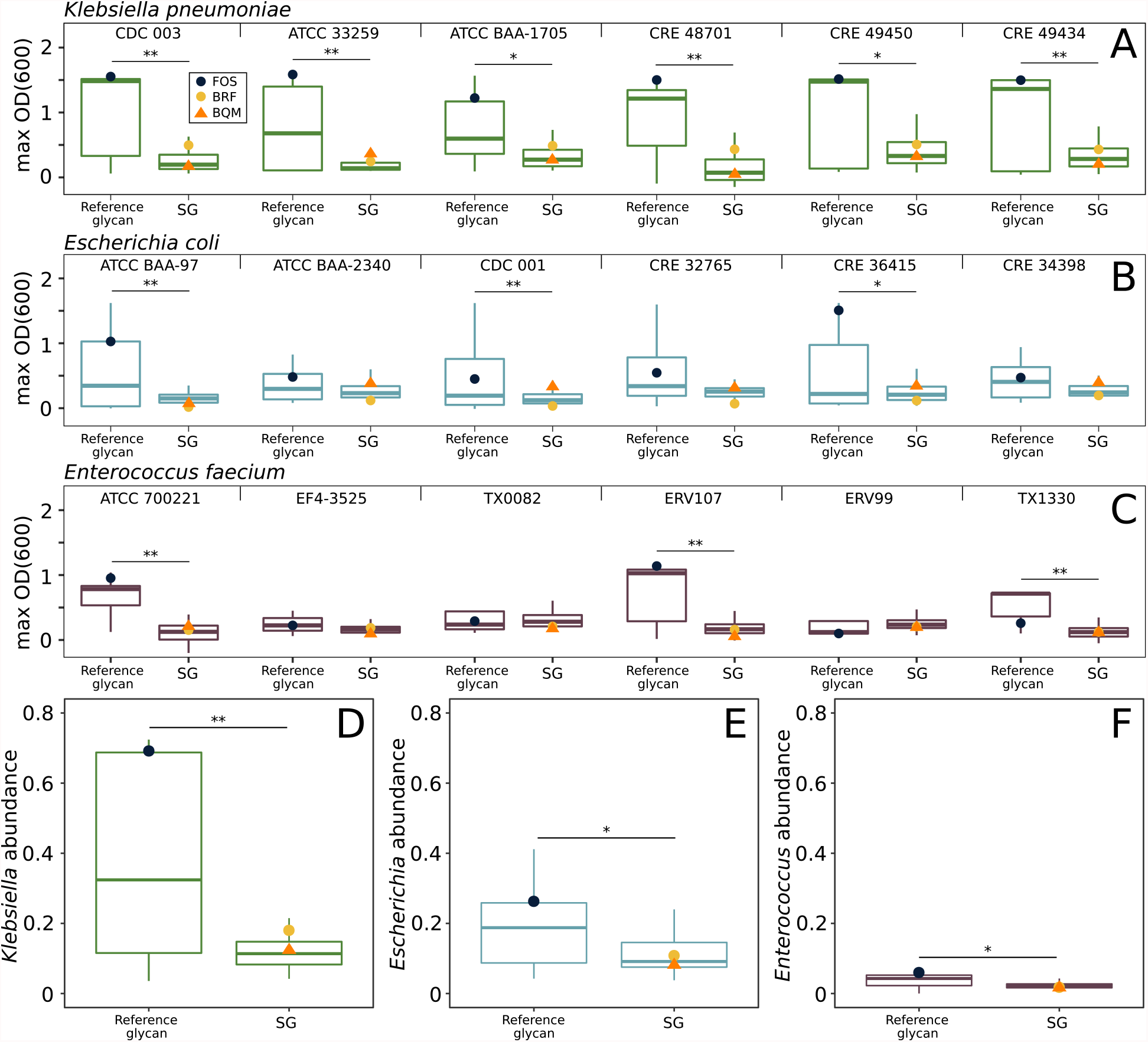
Enteric pathogen (A-C) growth in pure culture and (D-F) relative abundance in fecal communities grown on glycans. Six strains of (A) *Klebsiella pneumoniae* (B) *Escherichia coli*, or (C) *Enterococcus faecium* were cultured with 5 g l^-1^ of a single SG (n=148) or reference glycan (n=32) in CM3 medium. Strain names are shown above each plot; data is the mean maximum optical density (OD_600_) of triplicate cultures. Fecal communities from a healthy donor were OD_600_-normalized to contain 8% of (D) *K. pneumoniae* CDC 003, (E) *E. coli* CDC 001, or (F) *E. faecium* ATCC 700221 and cultured in triplicate with 5 g l^-1^ of an SG (n=45) or reference glycan (n=17) for 45 hours in MM29 medium. The relative abundances of the pathogens were quantified by 16S rRNA gene sequencing. Data points show FOS (indigo circle), BRF (yellow circle), or BQM (orange triangle) cultures. Box plots show median and interquartile range. Asterisks show significance (*p<0.05, **p<0.01) by two tailed Student’s t-test. FOS, fructo-oligosaccharides; SGs, Synthetic Glycans; OD_600_, optical density at 600 nm.

To assess changes in the relative abundances of model pathogens in gut microbiomes during glycan fermentation, we spiked a healthy donor fecal community with either CRE *K. pneumoniae*, CRE *E. coli*, or VRE *E. faecium*, cultured it in medium containing each SG or reference glycan (Supplementary Table 4C), and measured changes in relative abundances of the pathogen by 16S rRNA gene sequencing. The relative abundances of *K. pneumonia* (Fig 3D), *E. coli* (Fig 3E), and *E. faecium* (Fig 3F) were significantly lower after growth on SGs including BRF and BQM than on reference glycans, consistent with the SGs favoring growth of commensal taxa.

In contrast to the limited growth of pathogens on SGs relative to reference glycans, pure cultures of phylogenetically-diverse, gut commensals (Firmicutes, Bacteroidetes, and Actinobacteria) grown in defined medium often reached similar cell densities with either BRF, BQM, or glucose as the sole carbohydrate source (Supplementary Fig 4A-F). We measured a time series of BRF and BQM consumption by a healthy donor fecal community using high performance anion exchange chromatography with pulsed amperometric detection (HPAEC-PAD), which showed the fecal community fully consumed these SGs by progressively metabolizing fractions of increasing DP (Supplementary Fig 5A,B).

### Structural features of SGs

Based upon the assimilated data from the *ex vivo* experiments, we focused on BRF and BQM because they have different monosaccharide compositions, are well-fermented, enrich different, anti-inflammatory taxa associated with resolution of colitis, promote high taxonomic diversity, and are poor growth substrates for pathogen-associated genera. Prior to testing these compounds in mouse models, we performed a detailed characterization of their chemical structures. The chemical catalysis methods for synthesis of BRF and BQM (see Methods) yielded an ensemble of structures that were defined as a function of monosaccharide input, catalyst, and reaction conditions^19^. Multiplexed capillary gel electrophoresis with laser-induced fluorescence detection (xCGE-LIF) showed BRF (Fig 4A) and BQM (Fig 4B) have low monomer content (1.1% and 2.2%, respectively). We observed a series of discrete peaks indicating multiple structures at each DP below five, but the increased structural complexities at higher DPs exceeded resolution. Size exclusion chromatography of BRF (Fig 4C) and BQM (Fig 4D) confirmed their low monomer contents and showed similar average DPs of 11.3 and 10.9, respectively. Both compounds have a polydispersity index of 1.9 due to the range of molecular masses comprising each SG.

**Fig 4.**
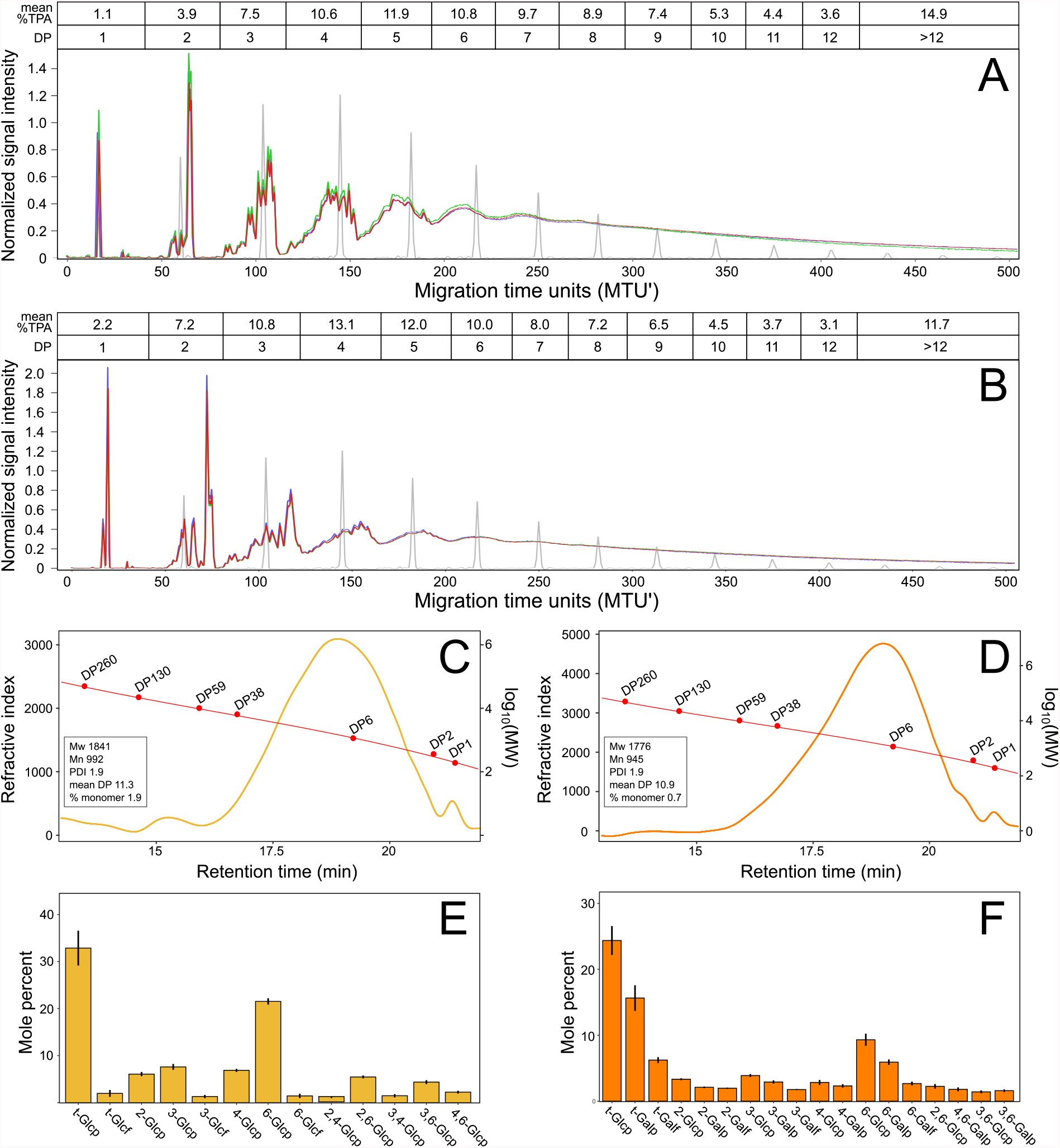
Structural features of SGs. xCGE-LIF analysis of the DP of (A) BRF and (B) BQM. Triplicate measurements of each SG (red, green, purple) and oligo-maltose standards (gray) are shown relative to normalized migration time units (MTU’). The mean percent TPA at each DP are above the plots. SEC chromatograms of (C) BRF and (D) BQM showing distributions relative to molecular weight (MW) standards with insets showing polymerization parameters. Refractive index in millivolts and molecular weights in Daltons. (E-F) Abundances (mole percent) of monosaccharides with different glycosidic linkages in (E) BRF and (F) BQM. Labels show all linkage types at greater than 1% mole percent with residues linked only at 1-position as terminal “t-” residues. Bars show means ±SD for two independent syntheses of each glycan analyzed in duplicate. SGs, Synthetic Glycans; xCGE-LIF, multiplexed capillary gel electrophoresis with laser-induced fluorescence detection; SD, standard deviation; TPA, total peak area; DP, degree of polymerization; SEC, size exclusion chromatography; Mn, number average molecular weight; Mw, weight average molecular weight; PDI, polydispersity index; Glcp, glucopyranose; Glcf, glucofuranose; Galp, galactopyranose; Galf, galactofuranose.

We performed linkage analysis of BRF (Fig 4E) and BQM (Fig 4F) to examine the abundances and types of glycosidic bonds. Both glycans contained mixtures of 1,2-1,3-, 1,4-, and 1,6-bonds with frequent branching, as greater than two linkages were present in 19% of monosaccharide subunits in BRF and 14% of subunits in BQM. We further compared the glycosidic linkages of BRF and BQM relative to reference glycans by two-dimensional nuclear magnetic resonance spectroscopy (2D-NMR). The 2D-NMR profiles of BRF versus pullulan (Supplementary Fig 6A) and BQM versus GOS (Supplementary Fig 6B) support that these SGs contain a greater diversity of glycosidic bonds with distinct stereo-and regiochemistries, likely contributing to why these SGs are poor growth substrates for pathogens (Fig 3) and are comparatively slowly fermented by commensals (Fig 1F,G).

### SGs demonstrate therapeutic potential in mouse models

We evaluated the therapeutic potential of BRF and BQM in mouse models of intestinal damage and disease. Initially, we examined effects of BRF or FOS in a mouse model of dextran sodium sulfate (DSS)-induced colitis (Fig 5A). BRF treatment reduced DSS-mediated weight loss (Fig 5A) and improved scores of diarrhea (Fig 5B) and endoscopy (Fig 5C). Colonic histology showed DSS induced lesions including epithelial erosion or mucosal ulceration, loss of colonic glands, and inflammation of the mucosa, which was reduced by BRF treatment relative to DSS alone (Fig 5D). In contrast, FOS treatment failed to reduce weight loss or improve scores of stool, endoscopy or histology relative to DSS treatment without glycan supplementation (Fig 5A-D).

**Fig 5.**
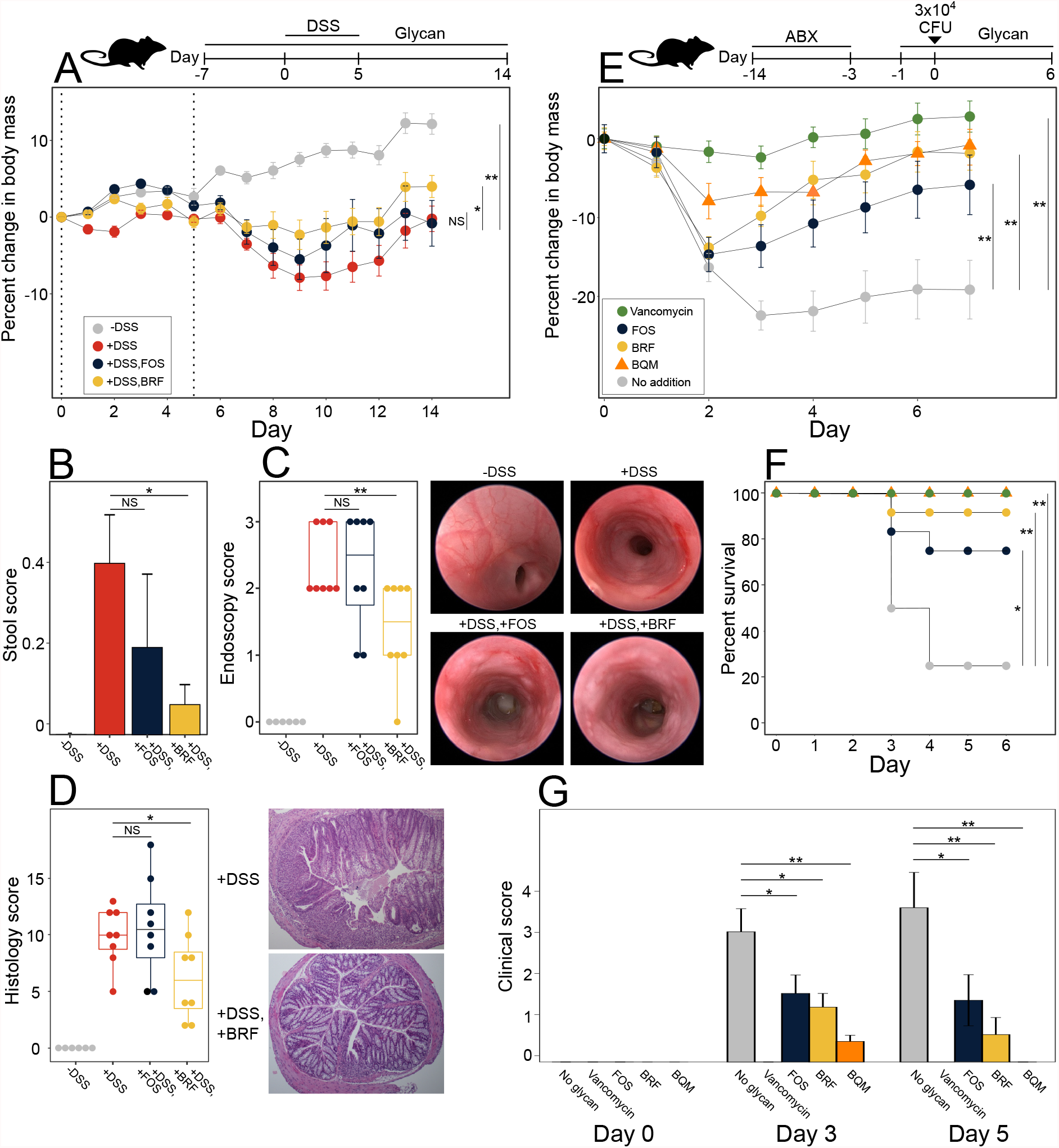
Glycan effects in mouse models of (A-D) DSS colitis and (E-G) *C. difficile* infection. (A-D) Mice were treated in drinking water with 2.5% DSS (days 0 to 5, dashed lines) and 1% (v/v) glycans (days -7 to 14), as appropriate. Treatment groups (8 animals per group): -DSS (gray), +DSS (red), +DSS, FOS (indigo), +DSS, BRF (yellow). Treatment group comparisons of (A) body weight, (B) stool score averaged over days 0 to 14, (C) day 14 endoscopy scores with representative images, and (D) day 14 histology scores with representative H&E stained distal colon micrographs. (E-G) Mice were treated with antibiotics (days -14 to -3), infected with *C. difficile* (day 0), and treated with 50 mg kg^-1^ vancomycin daily (days 0 to 4) or 1% (v/v) glycans in drinking water (days -1 to 6), as appropriate. Treatment groups (12 animals per group): no glycan (gray), vancomycin (green), FOS (indigo), BRF (yellow circles), and BQM (orange triangles). Treatment group comparisons of (E) body weight, (F) survival, and (G) clinical scores. (A, B, E, G) Data points show treatment group means ±SEM. (C, D) Box plots show median and interquartile range. (A, E) Statistics on body mass changes are based on area under the curve for all individual mice. Asterisks show significance (*p<0.05, **p<0.01) by (A, E, G) two tailed Student’s t-test, (B-D) Kruskal-Wallis followed by Dunn’s comparison test, (F) log-rank test. DSS, dextran sodium sulfate; FOS, fructo-oligosaccharides; ABX, antibiotics; CFU, colony forming units; SEM, standard error of the mean; NS, non-significant.

We also examined if BRF and BQM could alleviate pathologies associated with *C. difficile* infection. First, we confirmed that *C. difficile* grows poorly on BRF or BQM as a sole carbohydrate source in defined medium (Supplementary Fig 4G). We then tested if BRF, BQM, or FOS have *in vivo* efficacy in a *C. difficile* murine infection model (Fig 5E). Fecal metagenomics on day six after *C. difficile* infection revealed a divergence in fecal microbiome compositions across treatments (Supplementary Fig 7A) with the abundances of different genera elevated in each glycan treatment (Supplementary Fig 7B-E). Further, we found these microbiome changes translated into improved outcomes. Glycan treatment significantly reduced weight loss (Fig 5E) and increased survival (Fig 5F) relative to the no-glycan treatment. Whereas 3 of 12 mice survived in the no-glycan treatment, 11 of 12 mice survived BRF treatment and BQM treatment had 100% survival, similar to the vancomycin-treated controls (Fig 5F). In addition, all glycans improved clinical scores relative to the no-glycan treatment (Fig 5G). While FOS improved all metrics relative to the no-glycan treatment, BQM further reduced weight loss (Fig 5E), increased survivorship (Fig 5F), and improved clinical scores (Fig 5G) relative to FOS.

## Discussion

Perturbations to the taxonomic composition and metabolic output of the gut microbiome have been linked to numerous non-communicable and immune-related pathologies. Delivery of rationally-optimized, complex glycans is a promising option to drive the composition of the microbiota toward health-promoting states by leveraging taxa-specific differences in glycan metabolism, one of the dominant factors shaping gut microbiome composition^7^. Complex glycans such as SGs can alter gut microbiome composition because they resist cleavage by human-encoded enzymes, thus requiring the concerted action of bacterial carbohydrate-active enzymes (CAZymes) to break the glycosidic bonds^37, 38^. Only certain bacterial species in the gut microbiome possess the relevant CAZymes and metabolic capacity to utilize a given glycan, so different glycans promote the growth of specific taxonomic subsets of the microbiome^39-41^.

Our study demonstrates SGs are a novel, glycan-based modality to manipulate the properties of human gut communities with flexibility and precision. SGs are synthetic compounds with mixtures of glycosidic bonds that recapitulate features of the chemical complexity of dietary fiber, while avoiding challenges associated with natural fibers including geographic and seasonal variability of raw material sourcing. Moreover, SGs are well metabolized by polysaccharide-utilizing commensals, but not important pathogens. While still requiring validation in human trials, SGs could represent an antibiotic-sparing approach to control infection with no known mechanism of resistance.

Both of the SGs that we tested in mouse models shifted fecal community composition to elevate potentially anti-inflammatory taxa that are negatively associated with inflammatory bowel disease (IBD) including *Lachnospiraceae* (especially *Roseburia* and *Fusicatenibacter*), *Ruminiclostridium*, and *Parabacteroides*^31-33, 42^. As such, the SGs BQM and BRF could potentially alleviate altered microbiome compositions associated with mild and moderate inflammation, which is supported by how these SGs lessen pathologies associated with DSS colitis (Fig 5A-D) and *C. difficile* infection (Fig 5E-G) in mice. SG-mediated microbiome shifts could also be beneficial for other diseases. *Roseburia* is reduced in the microbiomes of colorectal cancer patients^43^. *Fusicatenibacter* and *Parabacteroides* are proposed to promote resistance to *C. difficile* infection^44, 45^ and *Parabacteroides* is associated with decreased hepatic steatosis^46^.

SGs extend the benefits of reference glycans and avoid many of the limitations of microbiome interventions based on live bacteria including dosing, engraftment, and inconsistent manufacturing. SGs could also be applied to promote growth and engraftment of live bacteria therapeutics. SGs selected for clinical development can be produced using controlled manufacturing processes and have been evaluated as Generally Regarded as Safe (GRAS) for their intended use in clinical research, allowing generation of proof of mechanism information by streamlined, human interventions.

Together, our results show SGs are chemically diverse glycans that effectuate novel taxonomy and metabolic shifts to gut communities and reduce symptoms of colitis in mouse models. Future work will further expand the potential of SGs in disease management by modulating the composition and output of the gut microbiome.

### Online Methods

#### Synthesis of BRF and BQM

BRF was synthesized by combining D-(+)-glucose (100.0 g, 555.1 mmol), Dowex® Marathon™ C (7.0 g, 5% dry weight ratio to glucose, 29% moisture content), and 30.0 mL DI water in a1000 mL 3-necked round bottom flask equipped with an overhead stirrer, thermocouple plug, and a short-path distillation head. The mixture was stirred continuously at 100 rpm using a glass stirring shaft equipped with a Teflon half-moon paddle. The reaction mixture was run at 130 °C for 4 h. To quench the reaction, 60 mL DI water was added to the reaction mixture. The Dowex resin was removed by vacuum filtration through a fritted-glass filter. The solution was then diluted to 25.0 °Bx and purified by precipitation. The solution was slowly poured into absolute ethanol to form a cloudy solution with a final water:ethanol ratio of 1:9 (v/v). The cloudy solution was then centrifuged at 4000 rpm for 2 h. The supernatant was removed, and the precipitate was collected and dissolved in water. The residual ethanol was removed under reduced pressure. The solution was frozen at -20 °C and lyophilized to yield the final product of white powder (60.5 g, 61% yield).

BQM was synthesized by combining D-(+)-glucose (50.0 g, 277.5 mmol), D-(+)- galactose (50.0 g, 277.5 mmol), Dowex® Marathon™ C (7.0 g, 5% dry weight ratio to monosaccharides, 29% moisture content), and 30.0 mL DI water in a 1000 mL 3-necked round bottom flask equipped with an overhead stirrer, thermocouple plug, and a short-path distillation head. The mixture was stirred continuously at 100 rpm using a glass stirring shaft equipped with a Teflon half-moon paddle. The reaction mixture was run at 130 °C for 4 h. To quench the reaction, 60 mL DI water was added to the reaction mixture. The Dowex resin was removed by vacuum filtration through a fritted-glass filter. The solution was then diluted to 25.0 °Bx and purified by precipitation. The solution was slowly poured into absolute ethanol to form a cloudy solution with a final water:ethanol ratio = 1 : 9 (v/v). The cloudy solution was then centrifuged at 4000 rpm for 2 h. The supernatant was removed, and the precipitate was collected and dissolved in water. The residual ethanol was removed under reduced pressure. The solution was frozen at -20 °C and lyophilized to yield the final product of white powder (57.2 g, 57% yield).

### Size Exclusion Chromatography (SEC)

Glycan samples (300 mg) were resuspended in 10 mL water, 0.2 mm filtered, and 10 μL of glycan solution was injected to an Agilent 1100 HPLC system with refractive index (RI) detector equipped with a guard column (Agilent PL aquagel-OH, 7.5 × 50 mm, 5 μm, PL1149-1530), and two SEC columns (2X Agilent PL aquagel-OH 20,PL1120-6520) in tandem connection. The sample was run with 0.1 M NaNO_3_ mobile phase, 28 min run time, 0.9 mL min^-1^ flow rate with the column and RI detector at 40 °C. The peaks of the sample were integrated and the weight-average molecular mass (M_w_), number-average molecular mass (M_n_), mean degree of polymerization (mean DP), and polydispersity index (PDI) were determined using Agilent Cirrus GPC/SEC software, version 3.4.2. The calibration curve was generated from polymer standard solutions (10 mg mL^-1^) of D-(+) Glucose Mp 180, Carbosynth Ltd Standard; Maltose Mp 342, Carbosynth Ltd Standard; Maltohexaose Mp 990, Carbosynth Ltd Standard; Nominal Mp 6100 Pullulan Standard, PSS # PPS-pul6k; Nominal Mp 9600 Pullulan Standard, PSS # PPS-pul10k; Nominal Mp 22000 Pullulan Standard, PSS # PPSpul22k; and Nominal Mp 43000 Pullulan Standard, PSS # PPSpul43k.

### xCGE-LIF Oligosaccharide Analysis

The DP distribution of SGs were determined using a glyXboxCE™ (glyXera GmbH, Magdeburg, Germany) based on multiplexed capillary-gel-electrophoresis with laser-induced fluorescence detection (xCGE-LIF)^47^. SG samples were prepared using a glyXprep kit for oligosaccharide analysis (KIT-glyX-OS.P-APTS-48-01, glyXera). Briefly, samples were diluted 100-fold with ultrapure water and labeled with fluorescent dye 8-aminopyrene-1,3,6-trisulfonic acid (APTS). Two μL of each SG sample, 2 μL APTS Labeling Solution, and 2 μL ReduX Solution were mixed thoroughly and incubated for 3 h at 37 °C. The labeling reaction was stopped by adding 150 μL Stopping Solution and the excess of unreacted APTS and salt were removed using glyXbeads for hydrophilic interaction chromatography solid phase extraction (HILIC-SPE). The samples were aliquoted to wells containing 200 μL glyXbead slurry and incubated for 5 min at ambient temperature for binding, followed by washing and elution steps.

The purified APTS-labeled glycans were analyzed on a glyXboxCE™ system (glyXera) equipped with a 50 cm 4-capillary array, filled with POP-7™ polymer (4333466 and 4363929, Thermo Fischer Scientific). The sample (1 μL) was mixed with 1 μL prediluted 1^st^ NormMiX (STD-glyX-1stN-100Rn-01, glyXera)^48^ internal standard to align migration times for migration time unit (MTU’) calculations. The mixture was combined with 9 μL glyXinject (C-glyXinj-1.6mL-01, glyXera) and subjected to xCGE-LIF analysis. The samples were electrokinetically injected and measured with a running voltage of 15 kV for 40 min. Data were analyzed with glyXtoolCE™ (glyXera) glycoanalysis software to perform migration time alignment, raw data smoothing, DP-range specific interval picking (summing up of the total measuring signal within the DP ranges after comparison with a reference oligo-maltose ladder), and normalization of total peak areas.

### Glycan linkage analysis

Permethylation was performed as previously described with slight modification^49^. The glycan sample was purified to a monosaccharide content < 1.0% using chromatography before permethylation. The glycan sample (500 μg) was dissolved in dimethyl sulfoxide (DMSO) for 30 min with gentle stirring. A freshly prepared sodium hydroxide suspension in DMSO was added, followed by a 10 min incubation. Iodomethane (100 μL) was added, followed by a 20 min incubation. A repeated round of sodium hydroxide and iodomethane treatment was performed for complete permethylation. The permethylated sample was extracted, washed with dichloromethane (DCM), and blow dried with nitrogen gas. The sample was hydrolyzed in 2M trifluoroacetic acid (TFA) for 2h, reduced overnight with sodium borodeuteride (10 mg mL^-1^ in 1M ammonia), and acetylated using acetic anhydride/TFA. The derivatized material was extracted, washed with DCM, and concentrated to 200 μL. Glycosyl linkage analysis was performed on an Agilent 7890A GC equipped with a 5975C MSD detector (EI mode with 70 eV), using a 30-meter RESTEK RTX®-2330 capillary column. The GC temperature program: 80°C for 2 min, a ramp of 30 °C min^-1^ to 170 °C, a ramp of 4 °C min^-1^ to 245 °C, and a final holding time of 5 min. The helium flow rate was 1 mL min^-1^, and the sample injection was 1 μL with a split ratio of 10:1.

### 2D HSQC NMR

Structures of SGs were compared to reference glycans: pullulan (P4516 Sigma), xylo-oligosaccharides (Bio Nutrition Life-oligo prebiotic fiber XOS), or galacto-oligosaccharides (DOMO Vivinal GOS). Glycans were lyophilized and a 20 mg sample was dissolved in 200 μL of deuterium oxide (D_2_O) with 0.1% (v/v) acetone as the internal standard. The solutions were placed into 3 mm NMR tubes and HSQC (heteronuclear single quantum coherence) spectra were recorded at 25 °C on a Bruker AVANCE III 500 MHz spectrometer equipped with a 5mm BBF-H-D-05 probe with Z-axis gradient using the Bruker program “HSQCETGP.hires2”. HSQC experiments were performed with 8 scans and a 1 second recycle delay. Each spectrum was acquired from 7.5 to 1.5 ppm in F2 (^1^H) with 1024 data points and 120 to 50 ppm in F1 (^13^C) with 256 data points. The resulting spectra were analyzed using the MestReNova software (version: 12.0.0-20080) from Mestrelab Research.

### Glycan consumption analysis

Consumption of SGs during fermentation by fecal cultures was measured at ProDigest (Ghent, Belgium) by high performance anion exchange chromatography with pulsed amperometric detection (HPAEC-PAD) using a ICS-3000 chromatograph (Dionex, Sunnyvale, CA, USA) equipped with a CarboPacPA20 column (Dionex), as described previously^50^. Sample preparation involved initial dilution of the sample with ultrapure water followed by deproteinization with acetonitrile (1:1), centrifugation (21,380 × g, 10 min) and filtration (0.2 μm PTFE, 13 mm syringe filter, VWR International) prior to injection (5 μL) into the column. Qualitative glycan fingerprints were obtained by plotting the detected signal (nC) against the elution time.

### Fecal sample collection

Informed consent was obtained from all donors before fecal sample collection. To collect samples for *ex vivo* cultivation, donors were instructed to place the stool collection bowl and holder under the toilet seat in the center rear of the toilet to collect the bowel movement into the sample collection unit. The sample was closed, placed in a collection bag, and stored on cold packs until processing into a fecal slurry. To prepare fecal slurries for culturing, samples were transferred into filtered blender bags (Interscience) and diluted using 1x phosphate-buffered saline (PBS) and glycerol to result in 20% fecal slurry (w/w) containing 15% glycerol (w/w). Diluted samples were homogenized in a lab blender (Interscience 032230), flash frozen in a dry ice/ethanol bath, and stored at -80°C. Mouse feces were placed into a 96-well plate that was kept on wet ice and then frozen at -80°C. Separate disposable forceps were used for each cage of animals to avoid cross-contamination.

### Microbial cultivation

Fecal communities were cultured in Mega Medium 29 (MM29) (Supplementary Table 5A) and single strains were grown in Clostridial Minimal 3 (CM3) medium (Supplementary Table 5B). All cultures were grown anaerobically at 37°C containing 5 g l^-1^ of the appropriate glycan as the sole carbohydrate source. Growth was measured in 60 μl cultures in 384 well plates (3860 Corning) sealed using a Breathe-Easy® sealing membrane (Z380059 Sigma). Cell densities at an optical density of 600 nm (OD_600_) and pH were measured using a Biotek synergy H1 multi-mode plate reader outfitted with a Biostack 4 plate stacker. Fecal cultures were grown in MM29 medium for 45 hours before DNA extraction for metagenomic or 16S rRNA gene sequencing. Gas production was measured continuously in 25 ml cultures fermenting different glycans in MM29 medium grown in 125 mL glass bottles sealed with an Ankom^*RF*^ gas production module (Ankom, Macedon NY, USA).

Pathogen growth in pure culture was measured for strains each of *K. pneumoniae, E. coli*, and *E. faecium* (Supplementary Table 4A) cultured anaerobically in CM3 medium supplemented with 5 g l^-1^ of each glycan (Supplementary Table 4B). To measure pathogen abundance in fecal cultures, fecal slurries were OD-normalized to contain 8% of each pathogen strain (Supplementary Table 4A), grown for 45 hours in MM29 medium supplemented with 5 g l^-1^ of each glycan (Supplementary Table 4C), and relative abundances were quantified by 16S rRNA gene sequencing.

Culture pH was measured by supplementing the medium with 2 μM BCECF [2,7-bis-(2-carboxyethyl)-5-(and -6)-carboxyfluorescein (C3411 Sigma) and the ratio of BCECF fluorescence at the pH-sensitive point (485 nm excitation; 540 nm emission) relative to the pH-insensitive isosbestic point (450 nm excitation; 540 nm emission) was measured^51^. The pH was calculated relative to a standard curve of media at known Ph by fitting a sigmoidal curve using four parameter logistic regression. The R package, phgrofit, was developed to extract physiological descriptors from the kinetic pH and OD_600_ curves (Supplementary Fig 1, https://github.com/Kaleido-Biosciences/phgrofit/ for methods and code). Glycans were clustered based on twelve fermentation parameters that were transformed into Z-scores by subtracting the mean across all glycans and dividing by the standard deviation. Hierarchical clustering analysis on these Z-score values was performed using the Manhattan distance metric and the complete agglomeration method; cluster number was selected based on the “elbow method” to minimize the within cluster variation (sum of squares distance).

### DNA sequencing and analysis

Fecal microbiomes were characterized by 16S rRNA gene or shotgun metagenomic sequencing (Diversigen, MN USA). DNA was extracted from human and mouse fecal samples using Qiagen DNeasy PowerSoil extraction plates, quantified using the Quant-iT PicoGreen dsDNA assay (Thermo Fisher), and stored at -80°C until analysis. 16S libraries were prepared by PCR amplification with the 515F/806R primer set and metagenomic libraries were prepared using the NexteraXT kit before sequencing on the Illumina platform. The average reads per sample for shotgun metagenomics was 0.5 million reads and for 16S metagenomics was 25 thousand reads.

Sequences of 16S rRNA genes were analyzed by UNOISE clustering^52^ and denoising of raw sequences followed by DADA2/RDP taxonomic calling at the genus or family level^53^. Taxa count tables from mouse and human shotgun metagenomic sequencing data were generated by the SHOGUN pipeline^54^ using a database including the first 20 strains per species in RefSeq v87. Metagenomic reads with ambiguous species matches were excluded from taxa counts to reduce spurious matches.

To cluster SGs based on taxonomic response using metagenomics data from *ex vivo c*ultures, the relative abundance of each species was averaged across triplicate cultures for each glycan. K-means clustering was performed based on species relative abundances differences defined by species-level mapping of sequencing reads. The number of clusters (K=8) was selected using the “elbow method” to minimize the within cluster variation (sum of squares distance) while also minimizing K. For ecological analysis, relative abundance was averaged across technical replicates for each species, and the *vegan* R package^55^ used to compute the Shannon diversity, Bray-Curtis dissimilarity, and multidimensional scaling measures and coordinates. Standard R functions were used for all other data preparation, clustering, and statistical analysis.

### Short-chain fatty acid (SCFA) quantification

Culture supernatants and SCFA standards in culture medium were derivatized by 3-Nitrophenylhydrazine hydrochloride (3NPH, N21804 Sigma) through *N*-(3-Dimethylaminopropyl)-*N*′-ethylcarbodiimide hydrochloride (EDC, Sigma E6383) coupling^56^. Culture media (10 μL) was mixed with 10 μL of 100 mM EDC in 50:50 acetonitrile/water (v/v) containing 7.5% pyridine and 10 μL of 100 mM 3-NPH in 50:50 acetonitrile/water (v/v), incubated at 45°C for 1h, and diluted with 30 μL of 50:50 acetonitrile/water (v/v). The matrix internal standard (IS) solution was prepared by derivatizing 25mM of ^13^C-butyric acid and ^2^D-propionic acid using 3NPH and mixing with saturated 9-Aminoacridine (9AA, Sigma 92817) in 50:50 methanol/water (v/v) at a 1:4 volume ratio.

SCFA reaction mixtures were mixed 1:1 with matrix IS solution, co-crystallized onto a MALDI HTS target plate by a TTP Mosquito liquid handling system, and dried at room temperature. A Bruker Daltonics ultrafleXtreme MALDI TOF/TOF mass spectrometer was used to acquire mass spectra in reflectron negative ion mode using 65% laser from laser attenuator and 2200-shots accumulation for each sample. Data was collected and processed using Bruker PharmaPulse 2.1 software. Peak area of derivatized butyrate (m/z 222.2) and propionate (m/z 208.2) were monitored and normalized with the peak area of corresponding internal standards (m/z 224.2 and 210.2, respectively). The concentration of butyrate and propionate in each sample was calculated using calibration standards.

### DSS colitis mouse model

C57Bl/6 mice (Taconic Biosciences) at 6-8 weeks of age were randomized into cages each containing a treatment group of eight mice at Biomodels LLC (Waltham MA, USA). Animals were fed diet 5053 (Lab Supply, Fort Worth TX, USA) throughout the study with water provided *ad libitum*. All animals were weighed daily and assessed visually for the presence of diarrhea and/or bloody stool. Mice with greater than 30% weight loss were euthanized. Colitis was induced by supplementing the drinking water with 2.5% dextran sodium sulfate (DSS) from days 0 to 5. Glycans were administered in the drinking water as 1% (v/v) filter-sterilized solutions from days -7 to 14.

Stool consistency for each mouse was scored daily (Supplementary Table 6A). Colitis in each mouse was assessed by video endoscopy under isoflurane anesthesia on day 14. During each endoscopic procedure images were recorded and colitis severity was scored by a blinded observer (Supplementary Table 6B). Histopathological changes in tissue samples were assessed by a board-certified veterinary pathologist at Inotiv (Missouri, USA). Tissue samples were prepared by fixation in formalin and longitudinal and cross sections of proximal and distal colon were processed, embedded in paraffin, sectioned at approximately 5-6 μm, mounted on glass slides, stained with hematoxylin and eosin (H&E) examined microscopically, and scored using a qualitative and semi-quantitative grading system (Supplementary Table 6C).

### *C. difficile* infection mouse model

C57Bl/6 mice were randomly allocated into treatment groups of 12 animals (3 animals per cage) at Trans Pharm Pre-clinical Solutions (Jackson MI, USA). Mice were fed Teklad Global Rodent Diet 2918 (Harlan) and water *ad libitum*. All mice received an antibiotic cocktail (0.5 mg ml^-1^ kanamycin, 0.044 mg ml^-1^ gentamicin, 1062 U ml^-1^ colistin, 0.27 mg ml^-1^ metronidazole, 0.16 mg ml^-1^ ciprofloxacin, 0.1 mg ml^-1^ ampicillin, 0.06 mg ml^-1^ vancomycin, 1% (v/v) glucose) in their drinking water from day -14 to day - 5 followed by a single oral dose of 10 mg kg^-1^ clindamycin on day -3. On day 0, mice received an oral gavage of 3×10^4^ viable *C. difficile* spores (strain VPI 10463 ATCC 43255). Mice in the vancomycin control group received oral gavage of 50 mg kg^-1^ vancomycin daily from days 0 to day 4. Glycans were administered in the drinking water as 1% (v/v) filter-sterilized solutions from day -1 to day 6. Glycan efficacy was assessed by daily measurements of animal survival, body weight, and a clinical score of disease severity (Supplementary Table 6D).

## Data Availability

DNA sequencing reads fecal samples have been deposited in the NCBI SRA and will be publicly available upon acceptance of the manuscript.

## Code Availability

The methods and code for the phgrofit R package to extract physiological descriptors from the kinetic pH and OD_600_ data is available at https://github.com/Kaleido-Biosciences/phgrofit/.

## Acknowledgements

We would like to thank the Kaleido Discovery and Clinical groups for supporting the experiments, S Reiling, T Yatsunenko, and U Binné for discussions, AMRI for SCFA analyses, and L Jung (PrecisionScientia) for critical reading and editing of the manuscript.

## Author Contributions

ACT, GVM, CML, and JEHV designed research. NB, CB, BM, JM, YM, LL, GL, CM, RK and ER performed research. ACT, NB, CB, JT, JL, and RK analyzed data. ACT, PJT, and JEHV wrote the paper.

