## Supplemental Figures for "Synthetic glycans that control gut microbiome structure mitigate colitis in mice"

### Supplementary Figures

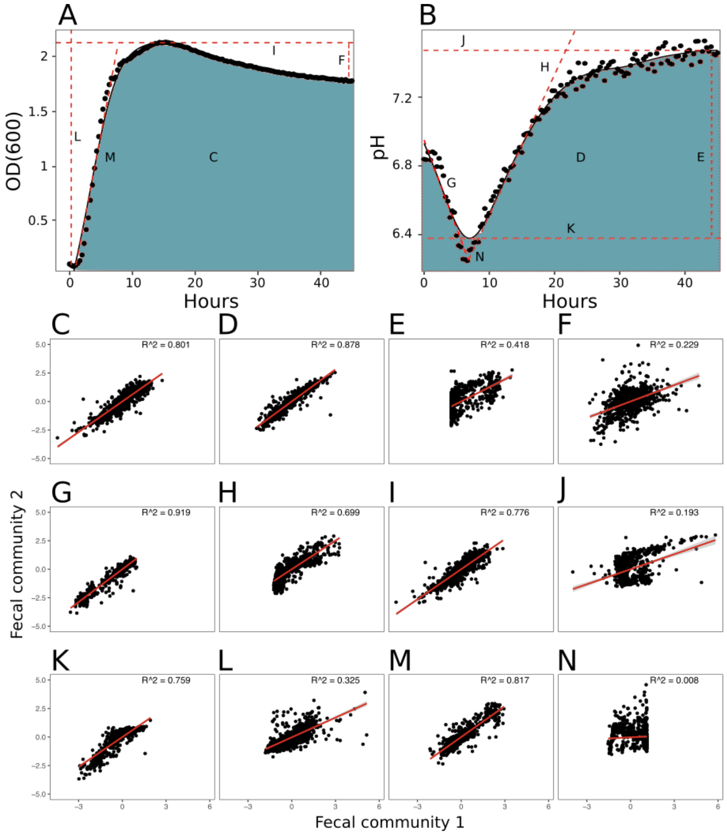


**Supplementary Fig 1** Method to measure glycan fermentation dynamics by extracting features from growth and pH curves, and its application to compare results between fecal communities from two healthy donors. Representative curves of (A) growth ($OD_{600}$) and (B) pH display features extracted from kinetic data fitted with smoothing splines. Letters in (A,B) correspond to features shown in panels C-N. (C-N) Comparison of fermentation features by fecal communities from two healthy donors grown in triplicate on 5 g $l^{-1}$ of each 653 SGs or 110 reference glycans as a sole carbohydrate source in MM29 medium. Correlation coefficients between two communities are shown. AUC of (C) growth and (D) pH data. (E) Difference between final and minimum pH. (F) Difference between maximum and final cell density. (G) Maximum acidification rate. (H) Maximum basification rate. (I) Maximum $OD_{600}$. (J) Maximum pH. (K) Minimum pH. (L) Growth lag measured where the tangent line to max growth rate intersects the starting cell density. (M) Maximum growth rate measured as maximum derivative of the growth curve. (N) Time of minimum pH. Values in C-N are shown are Z-scores calculated by subtracting the mean value for all compounds within a community and then dividing by the standard deviation of all compounds within a community. AUC was calculated using the trapezoidal rule. SG, Synthetic Glycan; $OD_{600}$, optical density at 600 nm; AUC, area under curve.

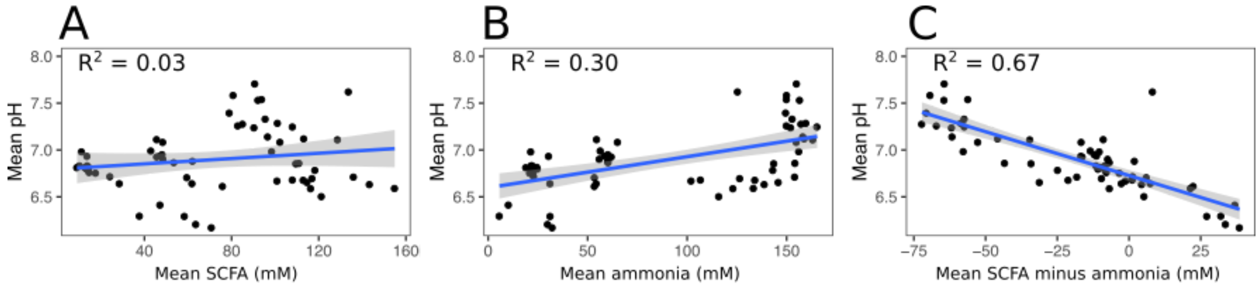


**Supplementary Fig 2** Correlations between glycan culture pH and (A) SCFA production, (B) ammonia production, and (C) relative production of SCFA and ammonia. Glycan fecal cultures were sampled for pH, total SCFA, and ammonia at 0,5,10,24, and 45h following inoculation. SCFA levels are the sum of acetate, propionate, and butyrate measured by gas chromatography with a flame ionization detector. Ammonia was measured by colorimetry (Abcam AB83360). Data points show means of triplicate cultures growing on 5 g $l^{-1}$ of randomly selected SGs in MM29 medium. SG, Synthetic Glycan; SCFA, short-chain fatty acid.

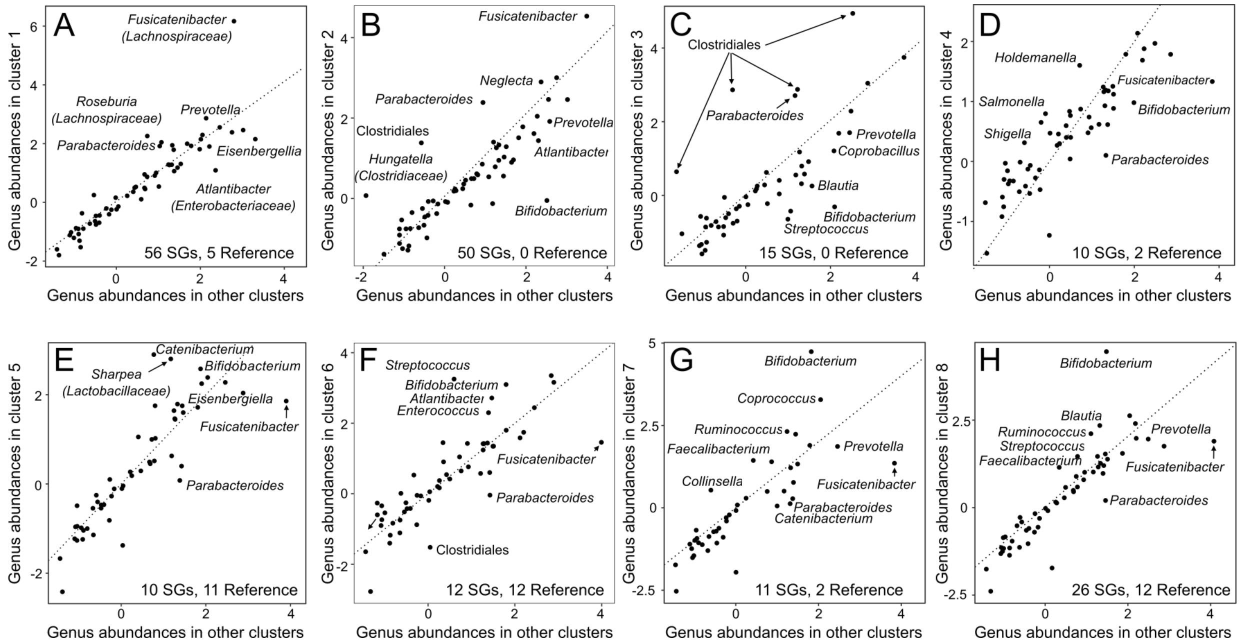


**Supplementary Fig 3** Glycan fermentation differentially shifts taxonomic compositions of fecal cultures. As shown in Fig 2G, glycan cultures were classified into eight K-means clusters based on species-level mapping of metagenomic sequencing reads with BRF in cluster 1 and BQM in cluster 2. Panels compare the abundance of each genus averaged across all glycans in the target cluster (y-axis) versus the abundance of that genus in the seven other clusters (x-axis). Genus abundances were calculated as $\log_2$(percent reads mapping to the genus in a glycan culture/percent reads mapping to the genus in the no-glycan control culture). SG, Synthetic Glycan.

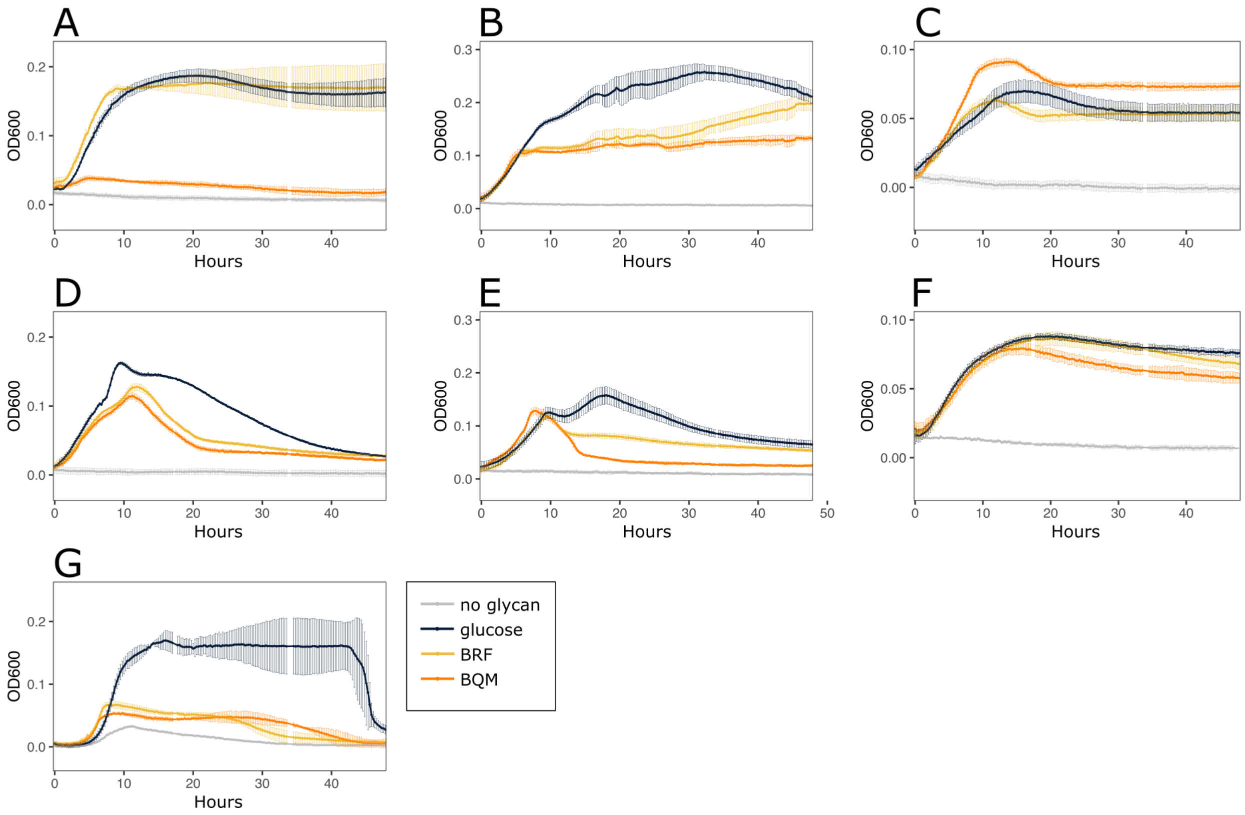


**Supplementary Fig 4** Growth of phylogenetically diverse gut bacteria on BRF, BQM or glucose as a sole carbohydrate source in defined medium. Cultures of (A) *Bifidobacterium longum* subsp. *longum* ATCC 15707, (B) *Bacteroides uniformis* ATCC 8492, (C) *Blautia hansenii* ATCC 27752, (D) *Parabacteroides distasonis* ATCC 8503, (E) *Parabacteroides merdae* DMSZ 19495, (F) *Collinsella aerofaciens* ATCC 35085, and (G) *Clostridium difficile* ATCC BAA-1382 were grown in CM3 medium with 5 g $l^{-1}$ of either BRF (yellow), BQM (orange), glucose (indigo) or without glycan supplementation (gray). Plots show mean cell density ($OD_{600}$) of triplicate cultures ±SD. SG, Synthetic Glycan; $OD_{600}$, optical density at 600 nm; SD, standard deviation.

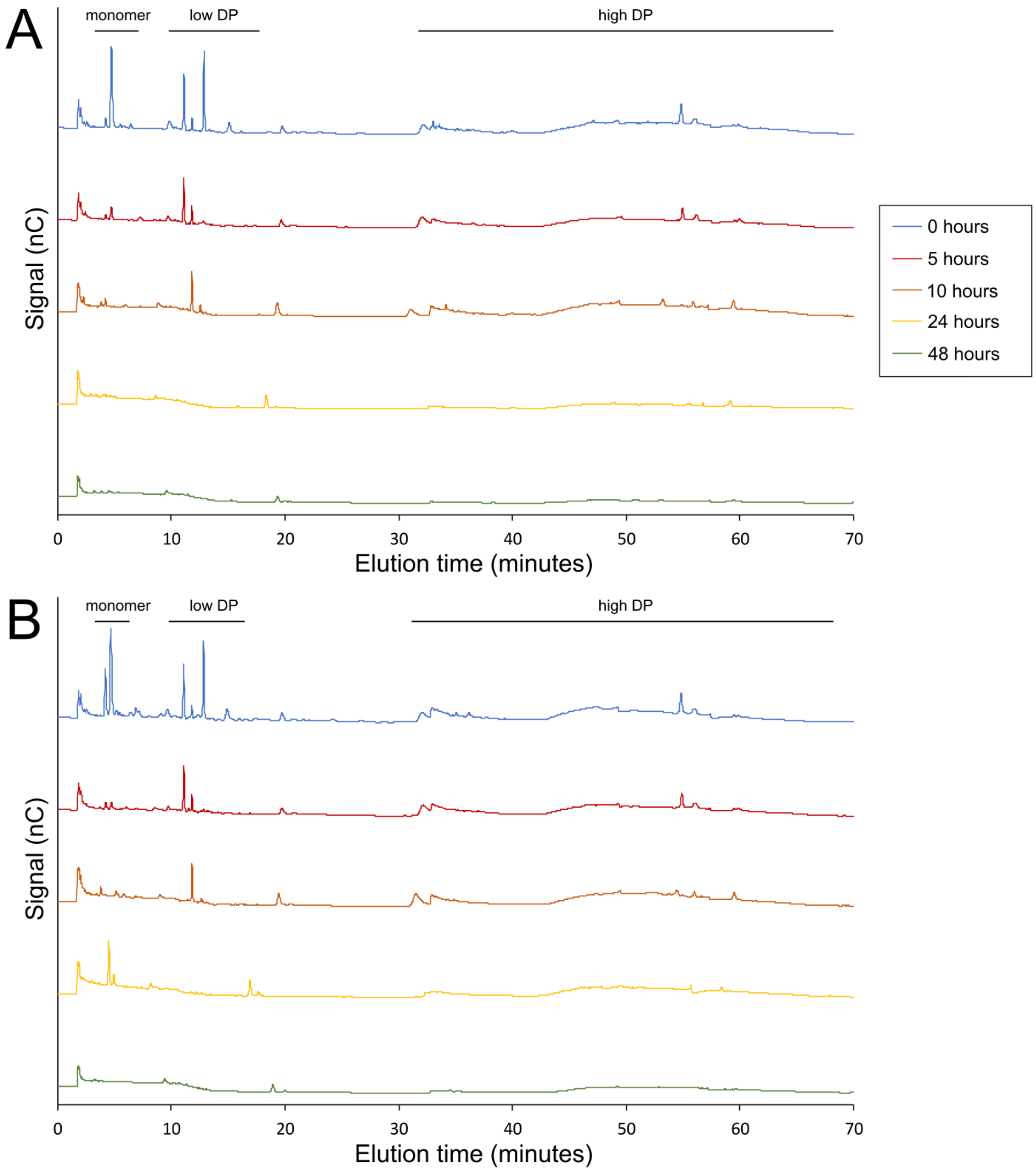


**Supplementary Fig 5** Chromatographic fingerprints showing consumption of (A) BRF and (B) BQM during fermentation by fecal cultures as measured by HPAEC-PAD. Cultures were grown at Prodigest (Ghent, Belgium) in SHIME medium (ProDigest) containing 5 g $L^{-1}$ SG for 48 hours at 37°C under anaerobic conditions. Glycan fingerprints after different incubation times (0, 5, 10, 24, 48 hours) are plotted as the detected signal (nC) versus the elution time (minutes). Elution periods corresponding to the glycan monomer, low DP fraction, and high DP fraction are shown above plots. HPAEC-PAD, high performance anion exchange chromatography with pulsed amperometric detection; nC, signal response, DP, degree of polymerization.

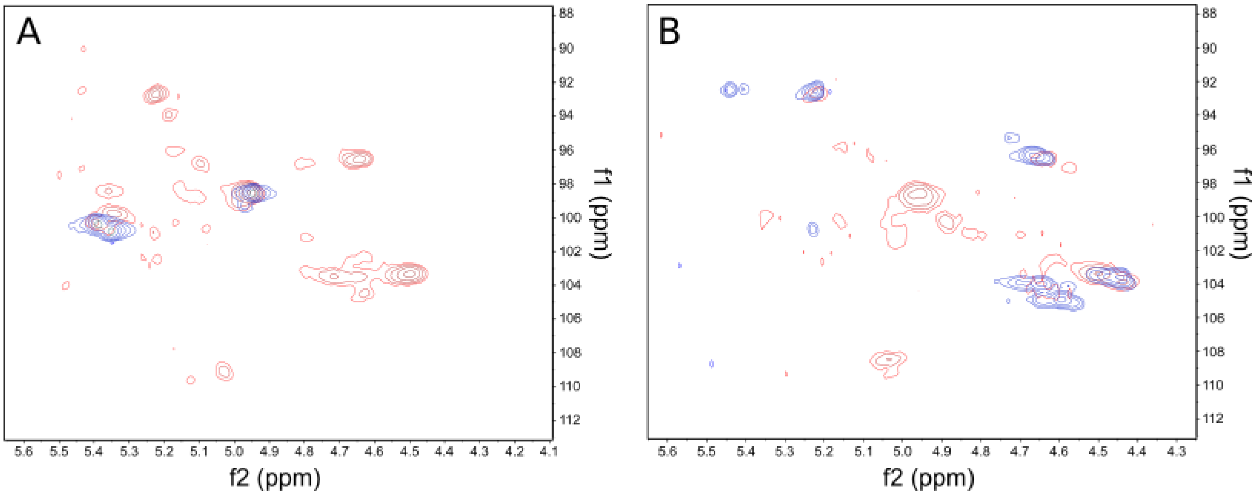


**Supplementary Fig 6** Comparison of SG and reference glycan glycosidic linkages. $^{1}H$-$^{13}C$ HSQC 2D-NMR spectra of the anomeric regions show that the stereochemistry and regiochemistry of glycosidic linkages are distinct and more numerous in SGs (red) versus reference glycans (blue). (A) BRF (100% glucose) versus pullulan. (B) BQM (glucose-galactose) versus GOS. Axes are $^{1}H$ along the f2 (horizontal) axis and $^{13}C$ along the f1 (vertical) axis. SG, Synthetic Glycan; HSQC, heteronuclear single quantum coherence; 2D-NMR, two dimensional nuclear magnetic resonance spectroscopy.

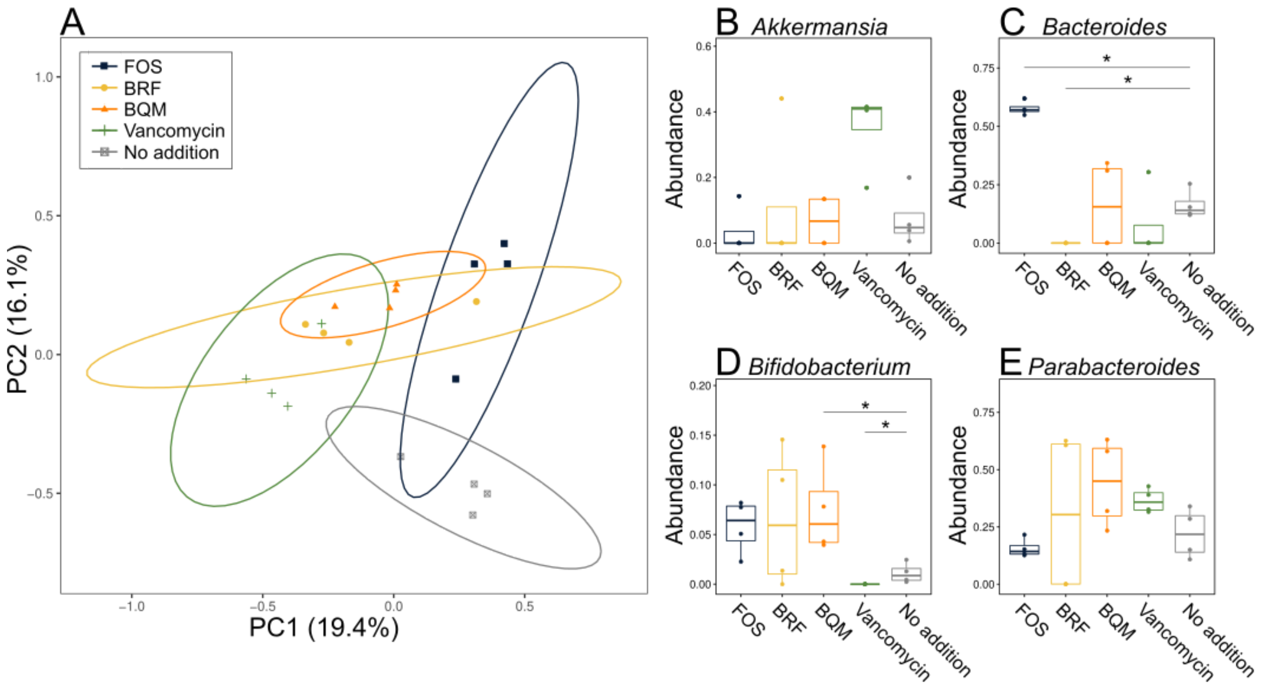


**Supplementary Fig 7** Glycan treatment changes the taxonomic composition of the mouse gut microbiome based on fecal 16S metagenomics in a *C. difficile* infection model. Treatment groups (12 mice per group): no glycan (gray), vancomycin (green), FOS (indigo), BRF (yellow) and BQM (orange). Metagenomics shows genus-level read mapping from fecal samples 6 days after *C. difficile* infection with each data point representing the mean of a cage with 3 mice. (A) PCoA of metagenomic data calculated using a matrix of Bray-Curtis dissimilarities shows divergent microbiome compositions across treatments. Axes show percent variance explained by each PC. Ellipses are 95% confidence intervals. (B-E) Comparison of relative abundances of genera based on percent mapped reads for (B) *Akkermansia*, (C) *Bacteroides*, (D) *Bifidobacterium*, and (E) *Parabacteroides*. (B-E) Box plots show median and interquartile range; asterisks show significance (*p<0.05) by Wilcoxon test. PCoA, principal coordinates analysis; PC, principal coordinate.
